# Proximate causes and risks of illegal grazing in Serengeti National Park: Perceptions of livestock keepers

**DOI:** 10.1101/2022.12.21.521527

**Authors:** Lameck Emmanuel Matungwa, Juma Joseph Kegamba, Alex Wilbard Kisingo, Masuruli Baker Masuruli

## Abstract

Ensuring the sustainability of protected areas for biodiversity conservation is a global issue that requires urgent attention for better conservation outcomes. Serengeti National Park (SNP) is a major tourist destination in Tanzania and offers diverse and spectacular wildlife attractions. The park is also a World Heritage Site, and there is no provision for legal grazing within the park. Understanding the proximate causes of illegal grazing of livestock in SNP and the perception of livestock keepers of the risks taken are critical to planning effective and sustainable mitigation strategies. This study used a semi-structured questionnaire to investigate the proximate causes of livestock grazing in the park and the perceptions of livestock keepers about the risks undertaken. We found that insufficient pasturage and water for cattle in the areas of stay, and the large number of cattle competing for common pasturage are perceived by the respondents as the proximate causes of livestock transgression into SNP. In addition to this, the free-range grazing system in Tanzania, the lack of land use plans, and climate change may be the main reasons for the decrease of pastures and the scarcity of water in the respondent’s areas and, therefore, lead to livestock transgression into SNP for supplementary forage and water. Furthermore, the results indicate that regardless of the number of cattle that the livestock keepers own, the majority fear being arrested inside the park by law enforcement patrols (wildlife rangers). However, most of Large Herders and Very Large Herders do not fear dangerous wild animals that might attack them or their livestock because they have different mechanisms of defense. We recommend that the responsible authorities consider revising the existing policy and promote more intensive livestock husbandry that encourages the management of pastures for livestock, emphasizes land use plans, and constructs farm dams and ponds for livestock keepers on the village land to increase retention by catchment and retention of rainy season water.

## Introduction

The anthropogenic impact of changes caused by human activity (agricultural or industrial activities) has resulted in the degradation of nearly a billion hectares of land globally [1]. Anthropogenic impact has been one of the main proximate causes of habitat loss [2], and the increasing isolation of protected areas in different parts of the world, including Africa, has affected the viability of wildlife populations and disrupted migration patterns [3]. The anthropogenic impact not only affects the presence and absence of species, but also influences the dominance of species in a given area [4]. One of the anthropogenic impacts that creates an increasing conservation challenge in wildlife-protected areas in Africa is illegal livestock grazing [5-7]. For example, livestock keepers in the western Serengeti have been demanding that the government degazette some of the protected areas for livestock keeping or legalize free access to pasture and watering points in Serengeti National Park (SNP), the Ikorongo and Grumeti Game Reserves (IGGR), and the Kijereshi Game Reserves [8-10]. Developing approaches to minimize livestock-wildlife interactions is vital for the continued existence of wildlife protected areas. Retaining expansive contiguous wilderness areas has been viewed as the most important global approach to conserving wildlife and species diversity [5, 10].

While the positive and negative impacts of grazing in wildlife-protected areas have been well-documented in the literature, there have been few studies in African [11, 12]. Livestock grazing in wildlife-protected areas has been found to increase plant species diversities [13] and promote seed dispersal [14]. However, numerous studies in African savannas have found that livestock grazing in protected areas affects the wildlife population negatively [15-17]. Livestock grazing in African protected areas has been implicated in water shortages, forage scarcity, and environmental degradation [17, 18]. Outside SNP, it was found that there is a clear deterioration of watersheds and river flows in areas where livestock grazing is carried out [19]. Similarly, it has been found that in a savanna ecosystem, a large number of cattle grazing in protected areas has resulted in the decline of the population sizes of large wild herbivores [5, 15, 16]. Livestock grazing in protected areas is also associated with the use of poison for retaliatory killing of wild carnivores to reduce livestock predation [20] and with the extraction of non-timber forest products for charcoal burning and building materials [7]. Furthermore, livestock grazing has been shown to alter wildlife habitat quality by damaging species composition and vegetation structure [21], reducing biodiversity and ecosystem functioning, and causing soil compaction, surface disruption, and reduced infiltration [22-24]. Brock and Green [22] singled out livestock grazing as a land use form that accelerates soil erosion and fragmentation of habitats compared to road development, housing, mining, and recreation.

The underlying root causes of the anthropogenic impact on the environment have been defined as “fundamental social processes, such as human population dynamics or agricultural policies, that underpin the proximate causes and either operate at the local level or have an indirect impact from the national or global level” [2]. In Tanzania, the root causes of livestock grazing in core protected areas have been identified as demographic factors, lack of appropriate policies, rapid growth of human and livestock populations, climate change, and poor land use plans [7, 10, 25]. Tanzania ranked third in the number of cattle in Africa after Ethiopia and Sudan, with 25 million cattle, 16.7 million goats, and 8 million sheep [26]. National Bureau of Statistics data for 2012/2013 [27] indicated that in Tanzania 50% of all households (4.6 million) kept livestock, with 82% and 23% in rural and urban areas, respectively. The current increasing trend and projections indicate in 2025 there will be 35 million cattle in Tanzania, with a large percentage kept free-range under transhumant pastoralism [28]. Therefore, wildlife-protected areas might be viewed as potential vacant land to absorb this growing pressure.

Proximate causes have also been defined as “human activities or immediate actions at the local level, such as agricultural expansion, that originate from intended land use and directly impact forest cover” [2]. For example, in the “W” Transboundary Biosphere Reserve in West Africa, the distance between protected areas and areas of residency for livestock keepers was perceived to be the proximate cause influencing livestock grazing in the protected area [29]. On the northern boundary of SNP and Kenya’s Mara Conservancy [30]—formerly known as the Maasai Mara National Reserve (MMNR)— the grazing land available outside the protected areas is small and insufficient, and the keeping of large herds of cattle (which is an implication of greater wealth), and the availability of grazing land were found to be the proximate causes of livestock transgression into protected areas. Butt [13] described this boundary crossing as a modern co-produced phenomenon and defined it as “incursion” within a complex political ecology framework. Other research [31] uses the less judgmental phrase “livestock transgression” to describe activities that are “not allowed.”

It should be noted that in East Africa (Burundi, Kenya, Rwanda, Tanzania, and Uganda), most of the 1776 nationally designated protected areas (PA), for wildlife and tourism, are protected through the fences and fines approach; therefore, unauthorized activities are associated with substantial risks [32]. Serengeti National Park (SNP) is a World Heritage Site and there is no provision for legal grazing in the park. Law enforcement patrols (wildlife rangers) check for illegal grazing. Suspected violators may be physically arrested—that is, stopped, charged, and taken into custody—or they may be charged and issued a summons to appear in court. Sometimes they are issued a fine and released, especially if a livestock keeper has money to pay the fine [33]. The livestock are seized until the fine is paid or the case is disposed in court, which sometimes results in livestock being confiscated. Most often the disposition is a fine or confiscation of livestock [34], rather than prison time, unless there are related charges such as possession of weapons, including local weapons such as machete, spears, and bow and arrow which can harm wild animals [33]. The amount of a fine is 100,000 TZS per single cow, which is multiplied by the number of cows found inside the park (Head of SNP Law Enforcement Department, Personal Verbal Comm. February 2017). Due to the risks of detection, most illegal activities are carried out at night [35]. For example, Knapp [35] found that when obtaining bushmeat from SNP, a poacher might face substantial risks, including personal injury, traveling long distances, and facing fines or a prison sentence if arrested. In the Mara Conservancy, in Kenya, livestock keepers were found to be at risk of arrest, attacks by wild animals on their livestock, and epizootic diseases such as sleeping sickness (spread by tsetse fly) and malignant catarrhal fever (spread by wildebeest) [13, 36]. The risks of grazing livestock in protected areas become much higher for livestock keepers who take livestock further inside the park, especially during the dry season, when pastures diminish in the periphery of the park [13]. Despite these risks, livestock keepers frequently transgress the borders of wildlife protected areas in search of alternative pastures after grazing areas in place of residency have been exhausted [37].

The increasing number of livestock entering protected areas threatens the sustainability of the wildlife and tourism industry in the eastern African nations. However, the integrity of protected areas can be maintained by preventing the pressure of livestock grazing and the encroachment of human settlement [37]. In 2013, local leaders around western Serengeti reported a massive internal migration of livestock keepers (and their cattle) from different regions of Tanzania settled in areas around SNP, and it was feared that serious damage to the ecosystem could result [38].

Increased livestock grazing pressure within SNP has raised major management concerns for the achievement of effective wildlife protection, as a large law enforcement effort is directed toward dealing with livestock transgression into the park (Head of SNP Law Enforcement Department, Personal Verbal Comm. February 2017). The key challenges in managing livestock on community land and the perceptions of livestock owners about the risks undertaken to graze livestock within SNP are poorly understood by conservation authorities. Thus, understanding the proximate causes of illicit livestock grazing in the park and the perception of livestock keepers about the risks taken is vital to better address this problem.

Specifically, we evaluated i) the proximate causes that influence livestock transgression into the park, and ii) the perception of livestock keepers toward the risks they take when grazing livestock in SNP. This research will provide information to the responsible authorities on how to assist livestock keepers with appropriate policies and plans to improve sustainable livestock management and reduce livestock transgression into SNP and other protected areas in the country. In addition to this, the research will broaden the knowledge of the research community and help resolve some of the challenges facing livestock keepers in the area.

## Materials and Methods

### The study area

The Serengeti-Mara Ecosystem (Serengeti Ecosystem) covers an area of 25,000 km^2^, and extends across the northwestern borders of Tanzania and Kenya (Fig. 1). The ecosystem is dominated by migratory wildebeest (*Connochaetes* spp.) that move across the border between the two countries. This ecosystem is made up of different protected areas with different administrative statuses [25, 39]. The Serengeti National Park (SNP), which covers 14,763 km^2^, is the core of the Serengeti Ecosystem [40] and is bordered to the north by <u>Kenya</u>, where it is contiguous with the Mara Conservancy in the Narok region. To the southeast of the park is the <u>Ngorongoro Conservation Area,</u> to the southwest lies the <u>Maswa Game Reserve,</u> to the west are the <u>Ikorongo</u> and <u>Grumeti Game Reserve</u>s, and to the northeast and east lies the <u>Loliondo Game Control Area.</u> The western part of the park, which is close to Lake Victoria, borders three districts, namely Bariadi and Busega in the Simiyu region and Bunda district in the Mara region

**Figure 1:**
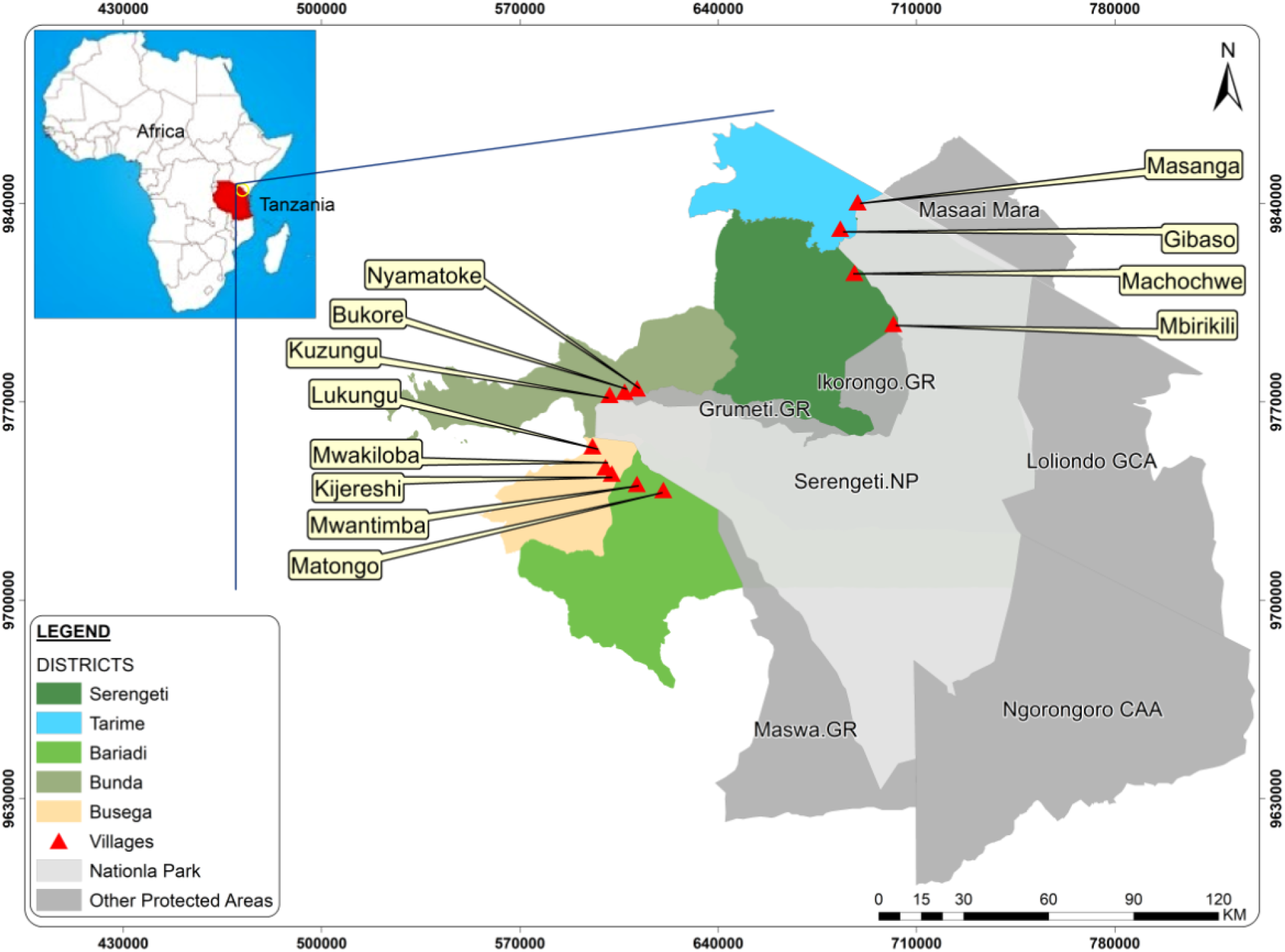
Map of the Serengeti Ecosystem showing the study area.

[40] (Fig. 1). This study was carried out in the western part of SNP, including 12 villages from five different districts (Serengeti, Tarime, Bunda, Busega, and Bariadi) that border the western part of the park (Fig. 1 and Table 1).

**Table 1:** Number of livestock keepers interviewed in each village

| Village name | District | Number of livestock keepers | Livestock Keepers Interviewed |
| --- | --- | --- | --- |
| Matongo | Bariadi | 394 | 58 |
| Mwantiba | Bariadi | 123 | 19 |
| Mbilikiri | Serengeti | 179 | 27 |
| Machochwe | Serengeti | 387 | 57 |
| Kijereshi | Busega | 143 | 36 |
| Mwakiloba | Busega | 181 | 27 |
| Lukungu | Busega | 139 | 20 |
| Gibaso | Tarime | 247 | 37 |
| Masanga | Tarime | 152 | 23 |
| Bukore | Bunda | 98 | 15 |
| Nyamatoke | Bunda | 81 | 12 |
| Kunzugu | Bunda | 123 | 19 |
| **Total** |  | **2347** | **350** |

The main economic activities of local communities living in western SNP are subsistence agriculture and livestock keeping [25, 41]. Cultivated crops include maize, cassava, millet, sorghum, and cotton, while livestock includes cattle, sheep, goats, and poultry. A household near western SNP owns an average of 17 animals with an average farm size of 0.9 to 3 ha [25]. Villages close to Lake Victoria are involved in small-scale and commercial fishing.

The Serengeti Ecosystem experiences a bimodal rainfall pattern with short rains from November to December and long rains from March to May; generally, the area receives a variation of rainfall from 500 mm / year in the south and east to 1150 mm / year in the north and western parts of the park [39].

### Data collection

We used semi-structured questionnaires [42] to collect data on the proximate causes influencing livestock keepers around SNP toward grazing cattle inside the park and the perception of those livestock keepers toward the risks undertaken. The main risks assessed in this paper refer to: 1) being arrested by wildlife rangers, and 2) encountering dangerous wild animals that can harm cattle caretakers or livestock. The survey was carried out from March to August 2017 in 12 villages that were selected from five districts adjacent to the park (Fig. 1), on the basis of preliminary information on livestock incursion incidents in SNP obtained from the Law Enforcement Warden (Head of SNP Law Enforcement Department, Personal Verbal Comm. February 2017). Before data collection, we obtained a research permit and presented an introductory letter from the College of African Wildlife Management, Mweka, to each District Executive Director (DED) (Ref. No. AC/S/15/Vol.II/42-46). From each district, we obtained another introductory letter, which was presented to the respective Village Executive Officer (VEO). We randomly (manually) selected households from a list of livestock keepers obtained from the VEO to conduct a questionnaire survey. Our focus was only on livestock keepers with cattle, and therefore we excluded those with only goats and sheep.

We categorized livestock keepers into four groups according to the number of cattle owned: Small Herders (1-20 cattle); Medium Herders (21-50); Large Herders (51-100) and Very Large Herders (>100). We pre-tested the questionnaire items following Hilton’s protocol [43] by administering it to 12 respondents from Matongo Village in Bariadi District and revised the questionnaire based on those pre-test results. For example, initially we thought livestock were restricted to graze and drink water within the village of stay, but that was not the case, as the pre-test results indicated that they are allowed to cross into other villages or wards within the district level depending on the availability of pastures and water. Then we revised our questionnaire and focused on the “*district*” as “*area of stay*” instead of a village.

Our questionnaire consisted of 20 items, including closed and open-ended questions (S1 Appendix), consistent with Hunter and Brehm’s method [44] that allowed respondents to express their own ideas and provide personal views. The first and second authors administered the questionnaire to the respondents. The questionnaire was divided into four sections: 1) a demographic profile of the respondent; 2) the knowledge of the respondent about the availability of pastures and water for livestock in their area of stay (district); 3) the experience of the respondent about grazing cattle inside SNP, specifically the risks undertaken; and 4) whether the respondent had any fear of being arrested or fear of problematic wild animals. The interviews began by seeking verbal approval from the respondents after they were informed of their right to withdraw from the interview at any stage and that whatever had been recorded would be redacted from the research records. In each interview, we asked the following questions: (i) Is there enough pasture for cattle in your areas of stay throughout the year? (ii) Is there enough water for the cattle in your areas of stay throughout the year? (iii) Have you ever taken cattle to graze in the park? (iv) Have you ever been arrested inside the park for grazing cattle? (Answers recorded ordinarily as 1, none; 2, once; and 3, multiple times) (v) Do you fear the risk of being arrested inside the park? And (vi) Do you fear the risks of dangerous wild animals that may attack you or your cattle inside the park?

The questionnaire was administered to the head of the selected household or any senior member of the household who was at least 18 years old, defining a household as a group of people living together, mostly with a single person self-identified as its head. The questionnaire was administered in the common language Kiswahili verbally and in writing, but when we encountered any language barrier (mainly with elders), we allowed the respondent to choose any member of the community for translation, and no one declined. Each respondent was assured that the information provided was only for study purposes and would be treated as confidential. Thus, as advised in Kaiser’s discussion of respondent confidentiality [45], each respondent was identified only by number. All selected households participated except a few where none of the senior members was available at the time a researcher visited their home, and replacements were selected from the nearby previously unselected household. Field observations were also conducted.

### Data analysis

We coded the data and entered them into SPSS 21 [46] before exporting them as a comma-separated values (CSV) file for analysis in R 3.6.2 [47]. We used generalized linear models (GLMs) repeatedly to assess which demographic variables (i.e., district, village, age, resident status (internal migrant or born in the village), and number of cattle) influenced the illicit grazing of livestock inside the park. We used GLMs with a binomial distribution error because the response variables were binary (1 = yes and 0 = no). Thus, we fitted six binomial GLMs to assess if:

1. There is enough pasture for cattle in the areas of stay throughout the year.
2. There is enough water for cattle in the areas of stay throughout the year.
3. Livestock keepers have ever taken cattle to graze inside the park.
4. Livestock keepers have ever been arrested with cattle inside the park.
5. Livestock keepers fear the risk of being arrested in the park.
6. Livestock owners fear the risks of dangerous wild animals inside the park.

After fitting the binomial GLMs, we used a likelihood ratio chi-square test (χ^2^) to select the best model terms using the *drop* function from *base-*R. This function assesses all possible models that can be fitted by performing a single model term deletion [48]. We then used the *ggplot2* R-package [49] for visualizations, and the plots were merged using the *patchwork* R-package [50]. All analyses were performed using R 3.6.2 [47].

## Results

We surveyed 350 households, i.e., 15% of the total households (∼2347) in the study villages (Table 1). All respondents (100%) were male within age groups of 20 to 35 years (20%, n=70), 36 to 55 years (20%, n=70), and over 55 years (60%, n=210). In total, the number of cattle owned consisted of Small Herders (22%, n=78), Medium Herders (42%, n=148), Large Herders (27%, n=95), and Very Large Herders (8%, n=29).

Bariadi district had the largest number of Very Large Herders (44.8%) followed by the Bunda and Serengeti districts, each with 20.7% (Fig. 2). A large majority (86%) of the respondents were born in the villages surveyed.

**Figure 2:**
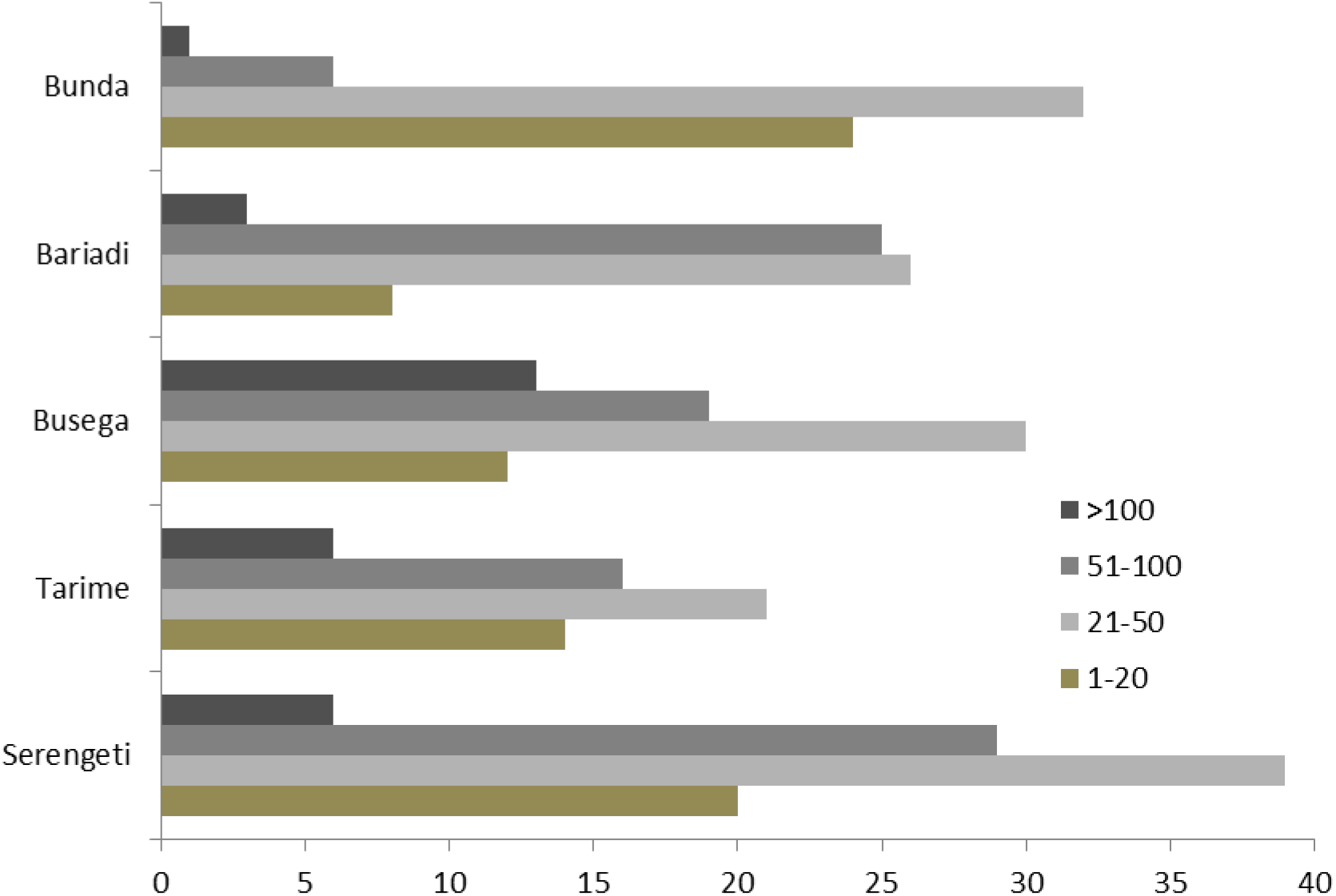
Number of cattle in categories owned by respondents from districts surveyed

### Availability of pasture in the area of stay

Respondents indicated that the pasturage in their areas of stay is not sufficient to accommodate livestock keepers with large herds. However, in the areas of stay (districts), more than half of the respondents in Bunda (56%) and Bariadi (53%) indicated that the pasturage is enough for the cattle they own, while 76% of the respondents in Serengeti district disagreed (Fig. 3a). Almost 92% of Small Herders responded that pasturage in their areas of stay are sufficient for their cattle throughout the year (Fig. 3b). However, 74% of Medium Herders, 79% of Large Herders, and 93% of Very Large Herders responded the opposite, saying that pasturage in their areas of stay are not enough for the cattle they own throughout the year.

**Figure 3a:**
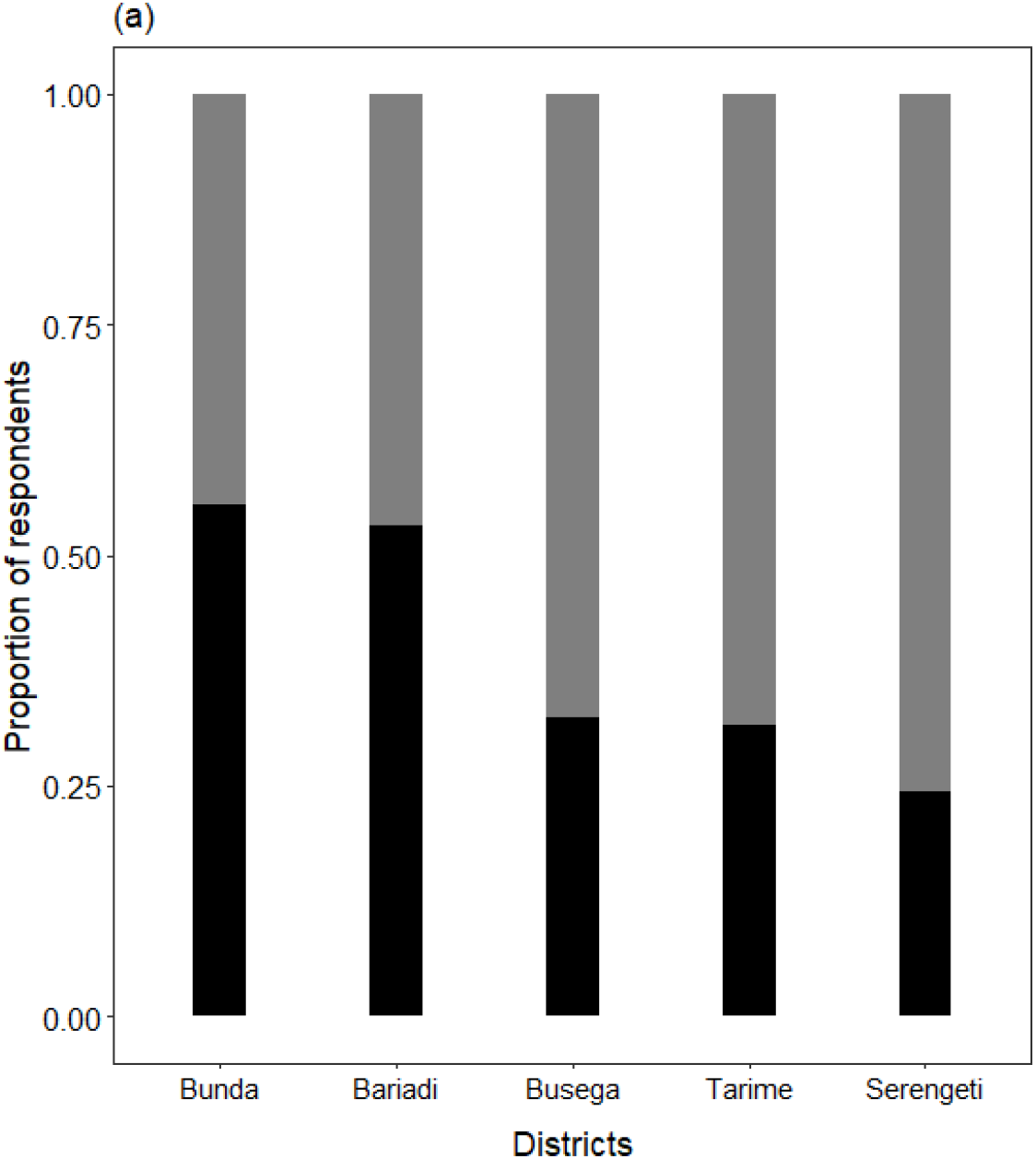
Sufficiency of grazing pastures per district

**Figure 3b:**
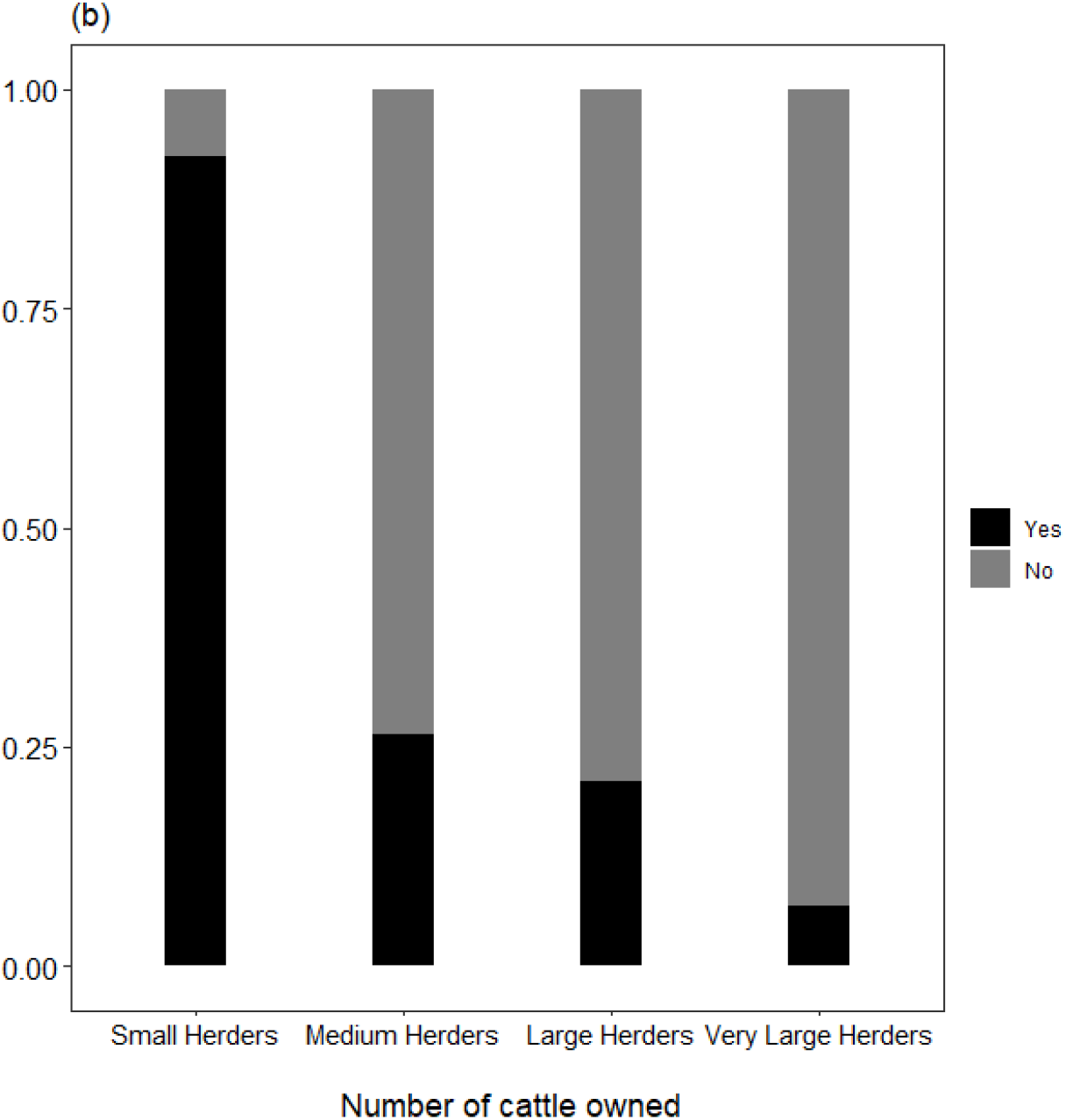
Sufficiency of pastures per number of cattle owned

The results showed that the area of residence (χ^2^ = 30.819, df= 4, p < 0.001) and the number of livestock owned by the respondents (χ^2^ = 133.605, df= 3, p < 0.001) were significant factors in predicting whether available pastures are sufficient for grazing livestock throughout the year. Other factors, including resident status (χ^2^ = 2, df= 1, p < 0.209), age (χ^2^= 2, df= 2, p < 0.324), and village (χ^2^= 8, df= 7, p < 0.359), did not influence the results.

### Availability of water for livestock in areas of stay

Similar to the responses on pasture availability, most of the respondents in Busega, Tarime, and Serengeti districts perceived that the availability of water in their areas of stay (districts) was insufficient to accommodate livestock keepers with large herds (Fig. 4a). However, almost half of the respondents in Bunda (53%) and Bariadi (50%) perceived that water is not a problem for the cattle they own. Furthermore, 72% of Small Herders responded that water for livestock in their areas of stay is sufficient throughout the year, while 97% of Very Large Herders disagreed (Fig. 4b). The results showed that the district where the respondent resides (χ^2^ = 15.190, df= 4, p < 0.0043) and the number of cattle owned by the respondents (χ^2^ = 57.439, df= 3, p < 0.001) were significant predictors determining the perception of whether the available water in the village was sufficient for the livestock. Other factors, including resident status (χ^2^ = 0.67, df= 1, p < 0.413), age (χ^2^ = 6.499, df= 2, p < 0.06), and village (χ^2^ = 8, df= 7, p < 0.359) did not significantly influence the results.

**Figure 4a:**
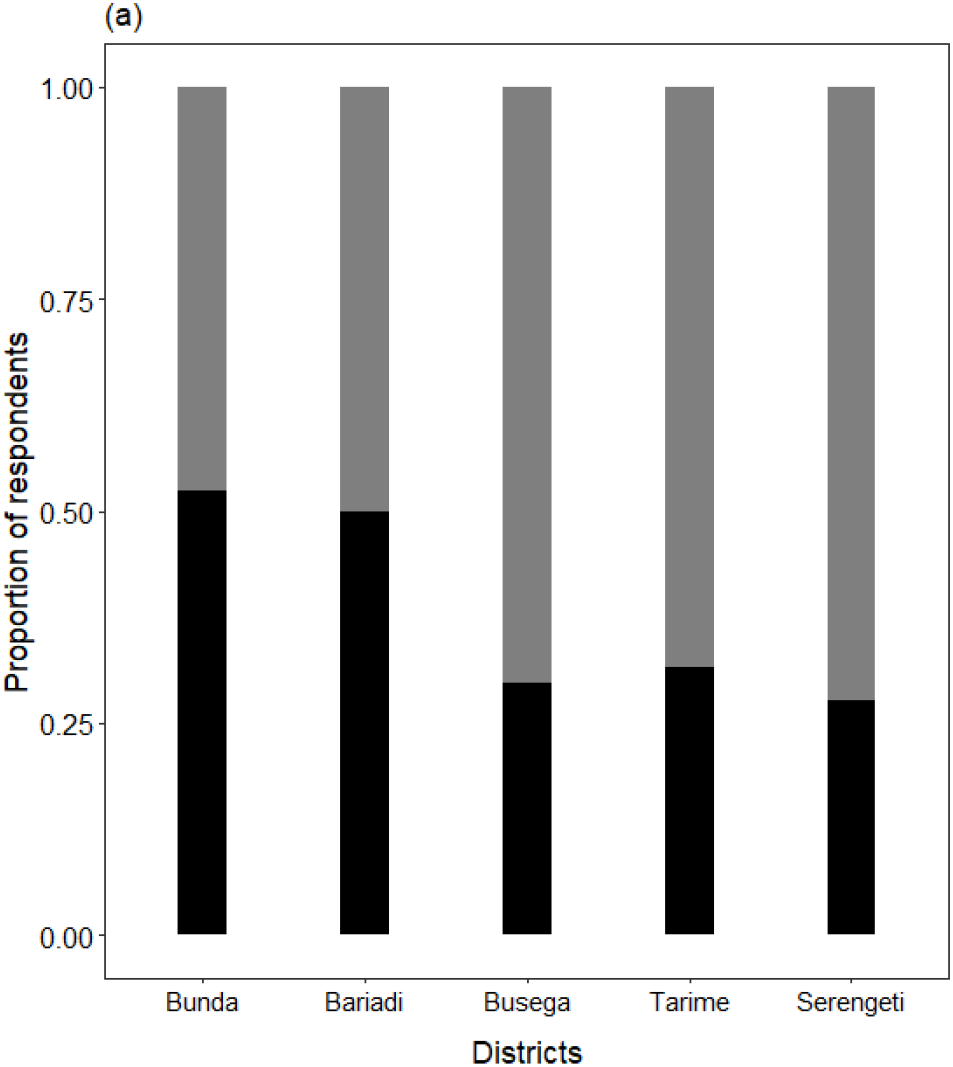
Sufficiency of water availability per district

**Figure 4b:**
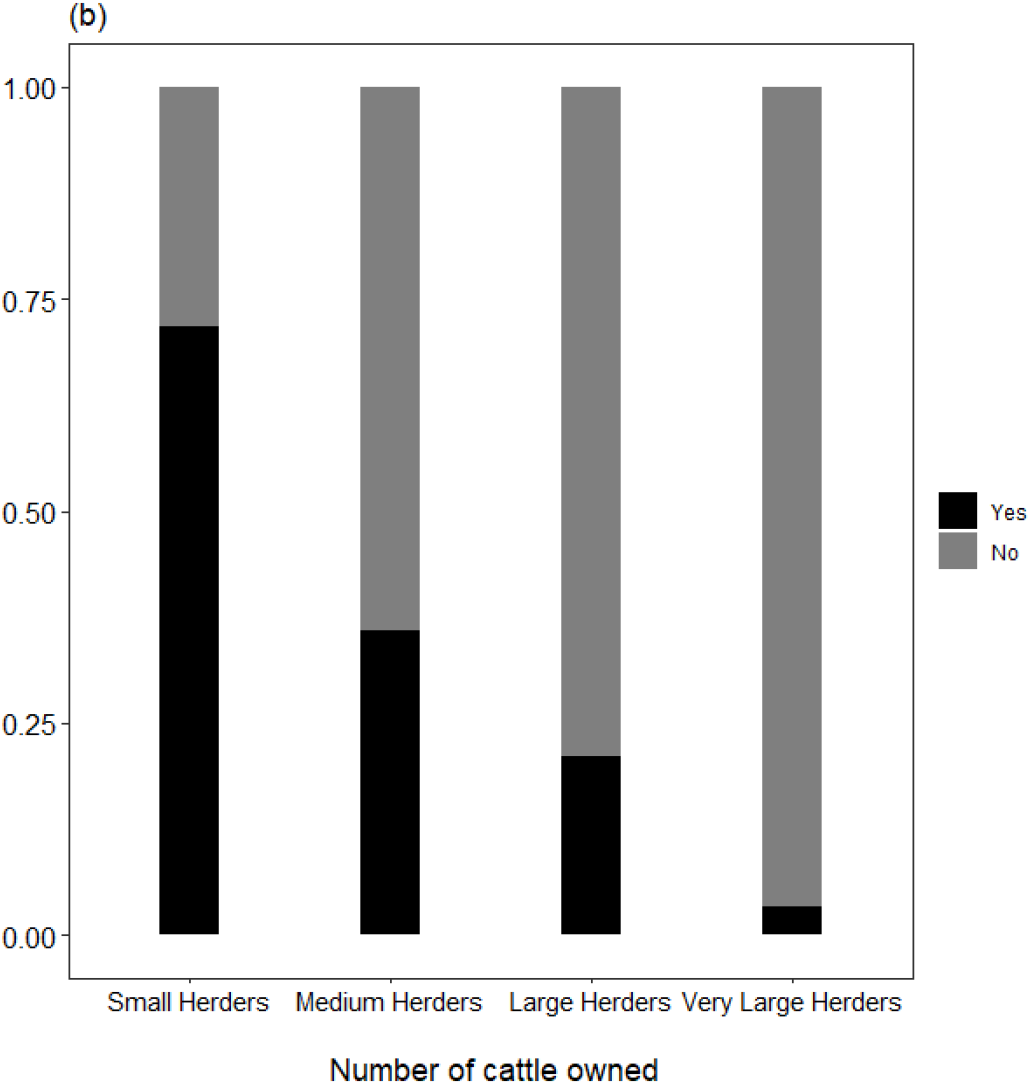
Sufficiency of water availability per number of cattle owned

### Grazing cattle and arrests in SNP

Most Large Herders (94%) admitted that, although illegal, they have taken livestock into the park for supplemental forage, and all Very Large Herders (100%) admitted that they have grazed livestock in SNP. However, 82% of Small Herders maintained that they have never grazed livestock in the park (Fig. 5a). Similarly, all Very Large Herders (100%) admitted that they have been arrested for grazing livestock inside SNP, while only 5% of Small Herders admitted having been arrested (Fig. 5b). Furthermore, the majority of Very Large Herders (83%) responded that they have been arrested multiple times in the park, while only 3% of Small Herders admitted to a single arrest, and none admitted to multiple arrests (Fig. 5c).

**Figure 5a:**
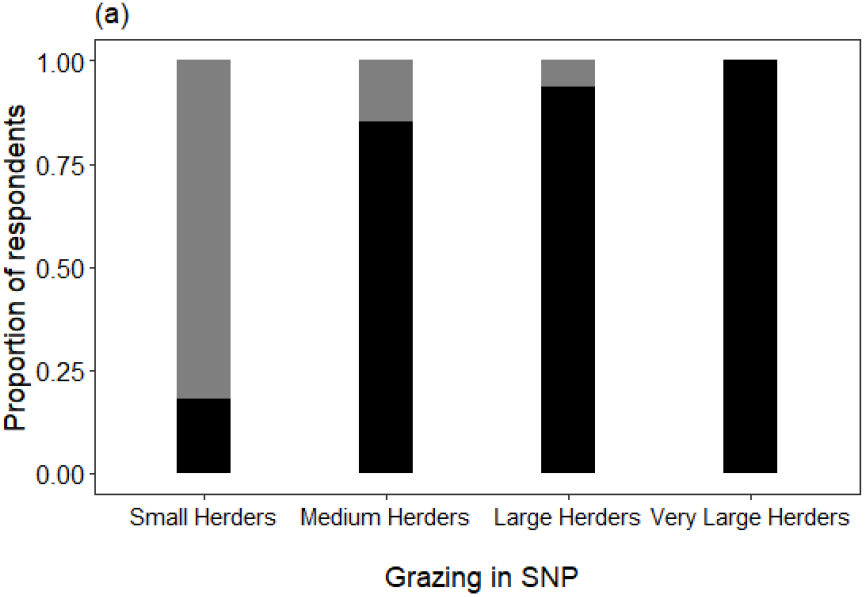
Proportion of herd size for respondents who have taken cattle to graze within the SNP.

**Figure 5b:**
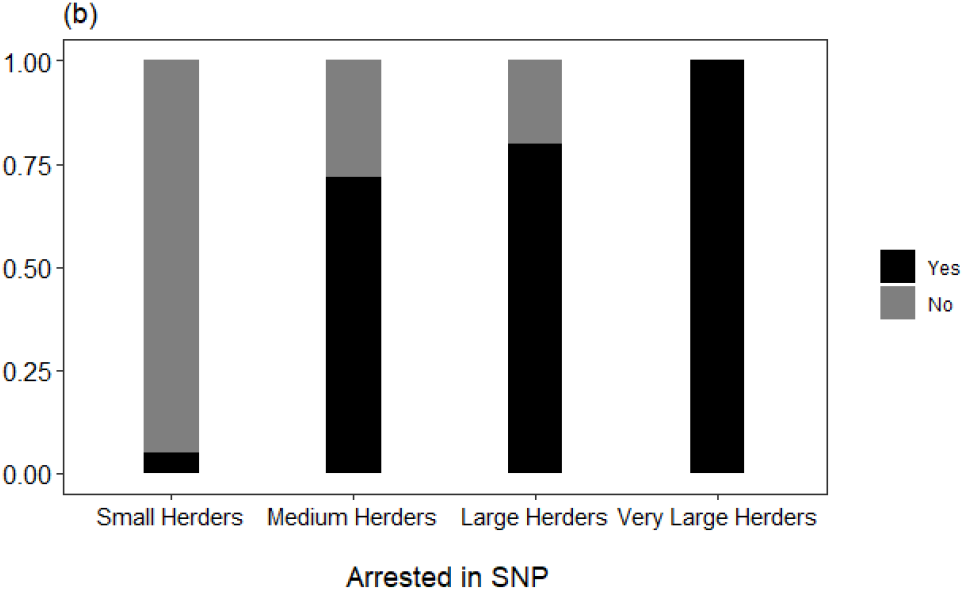
Proportion of herd size for respondents who have been arrested for grazing cattle in SNP.

**Figure 5c:**
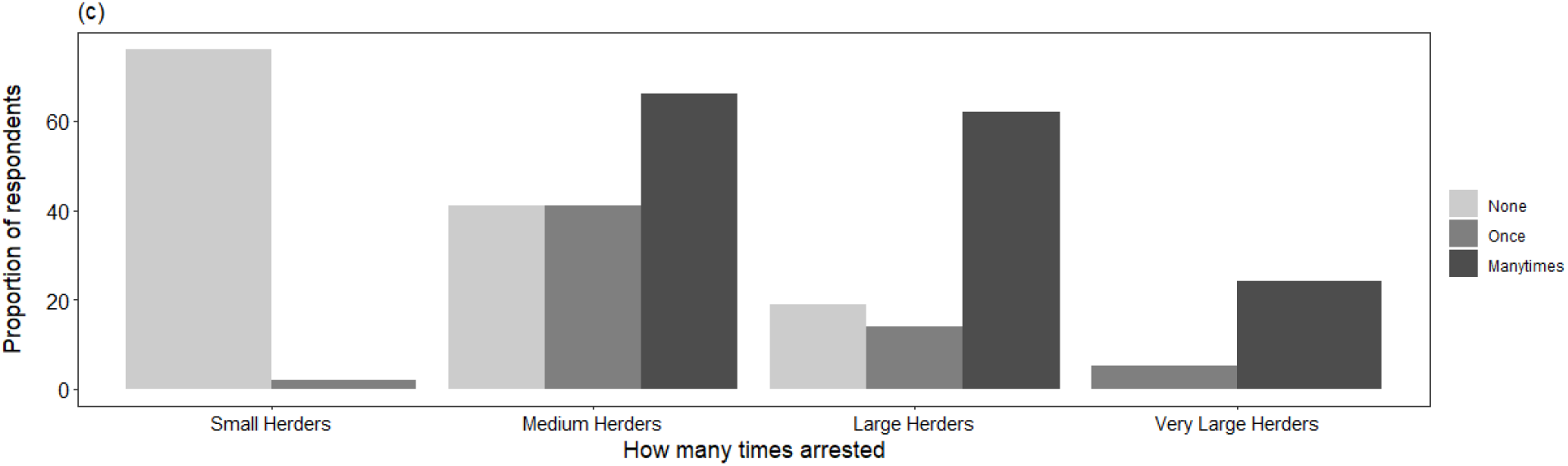
Frequency of arrests for respondents who have grazed cattle within the SNP

The results showed that the number of cattle the respondents own (χ^2^ = 140.759, df= 3, p < 0.001) was a significant factor in predicting the admission of whether the respondents have ever taken their livestock for grazing in the park, and also a significant factor in predicting the admission of whether a respondent has ever been arrested for grazing livestock inside the park (χ^2^ = 199.862, df= 3, p < 0.001). Other factors did not influence the results.

Field observations documented examples of routes used to access SNP (Figs. 6a-b). A total of 20 access routes were observed within 25 km along the boundary between SNP and the Bariadi district. Additionally, 12 access routes were observed within 25 km along the boundary between SNP and the Serengeti district.

**Figures 6a-b:**
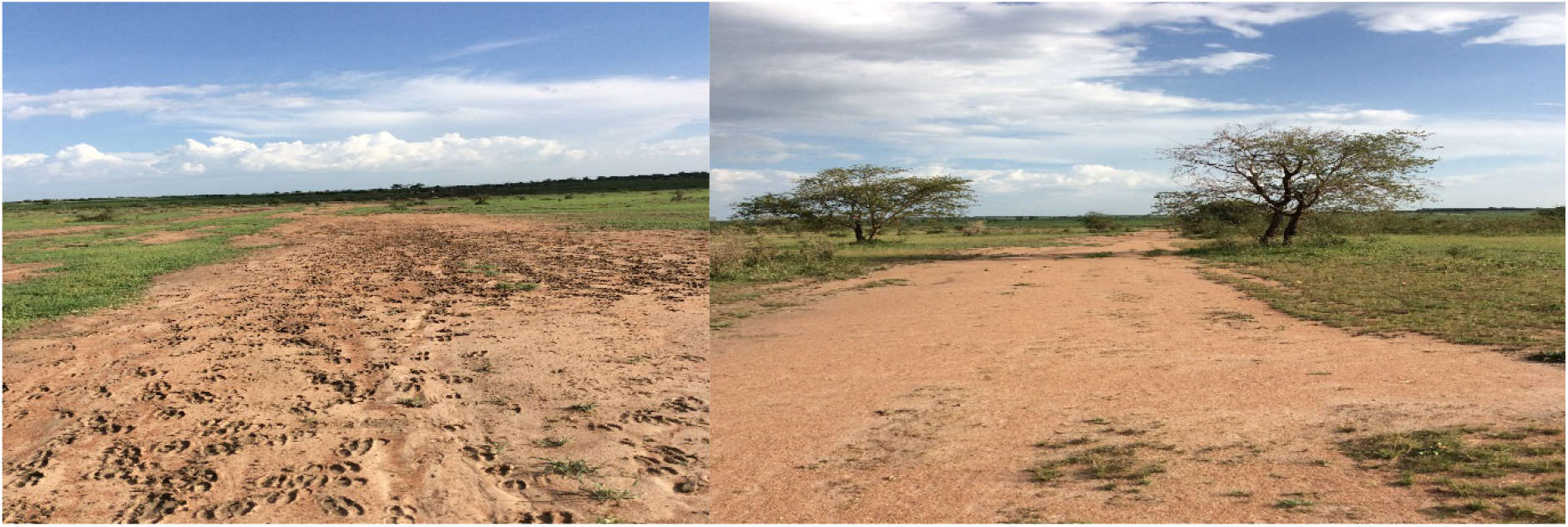
Observed access routes of livestock from villages to the park (Photo by the authors 2017)

### Perception of the risks taken by grazing cattle in SNP

Most of the respondents indicated that they almost equally fear being arrested inside SNP by wildlife rangers regardless of the number of cattle they own (Fig. 7a). However, 75% of Small Herders indicated that they fear dangerous animals, while only 17% of Very Larger Herders indicated this fear (Fig. 7b). The results showed that no factor (district of residence, age, residence status, or number of cattle owned) predicted a perceived fear of arrest, but the number of cattle the respondents own (χ^2^ = 22, df= 3, p < 0.001) was a significant factor in predicting the perceived fear of dangerous animals by the respondents when taking their livestock to SNP. Most of the respondents who said they do not fear dangerous animals acknowledged that they took a lot of risks, especially grazing their cattle at night in the park, and explained that they have a lot of experience and mechanisms for protecting their cattle, especially against carnivores, as cited in the following from two respondents (translated from Kiswahili):

**Figure 7a:**
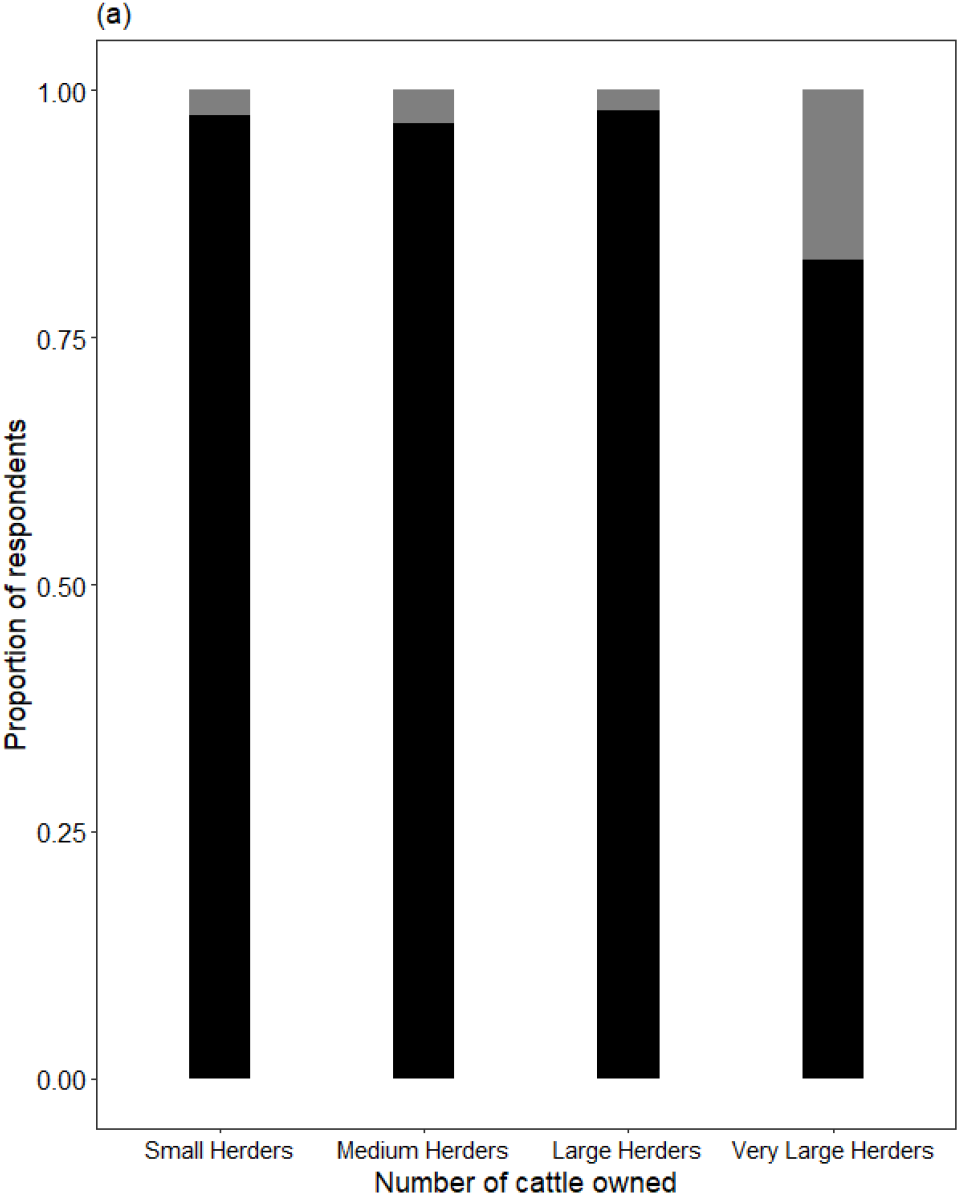
Respondent’s fear of arrest (for grazing cattle inside the park).

**Figure 7b:**
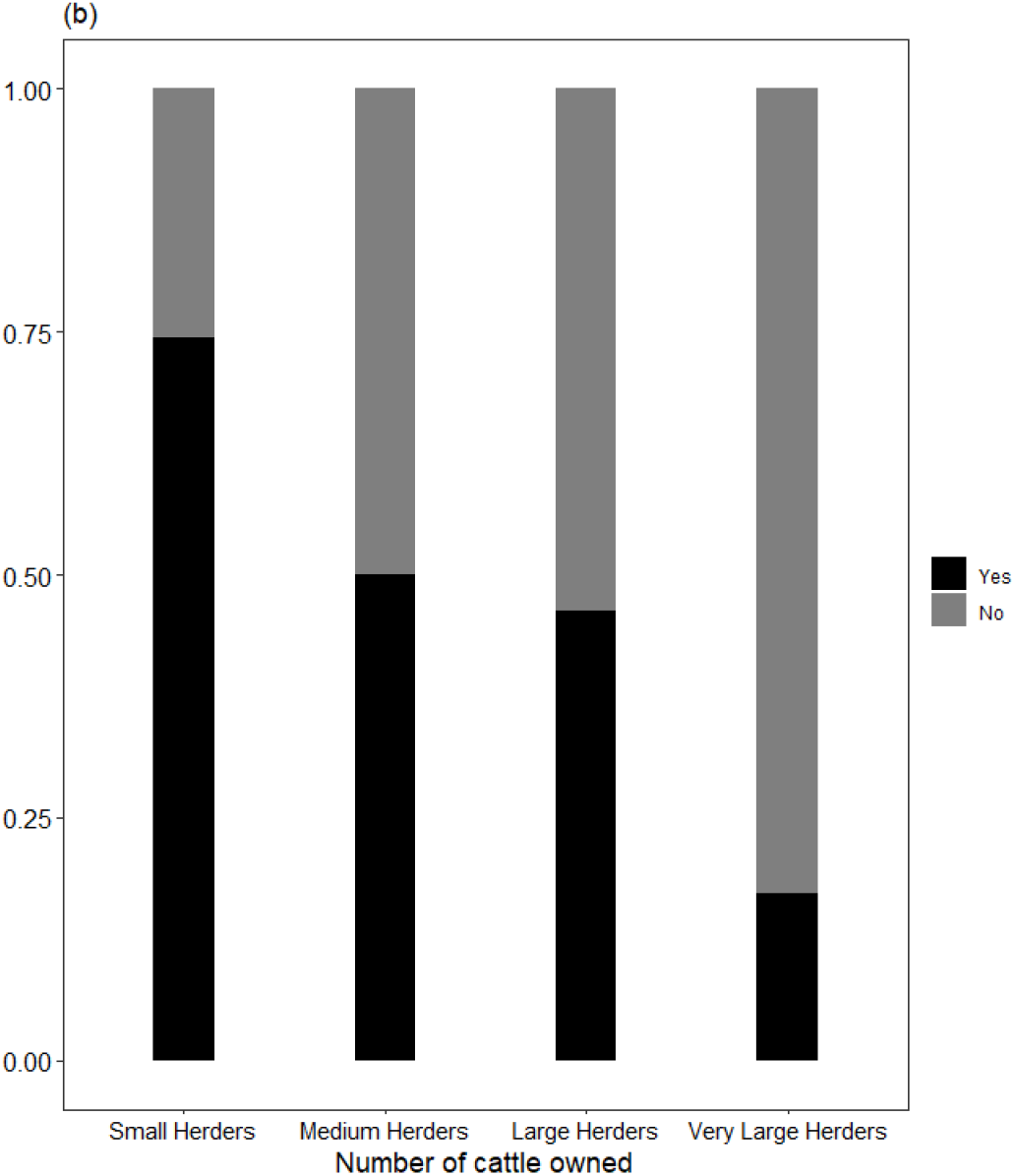
The fear of dangerous animals in the park of the respondent.

#### Respondent 1

“You have to choose—especially in the dry season—whether you lose most of your cattle as they starve in the village or whether you will sacrifice some of them when you take them to the park at night. We come across hyenas and lions, but you have to be strong to defend your cattle.”

#### Respondent 2

“Wild animals are scared of human being, and we use powerful torches at night to scare the most notorious ones like lions and hyenas. If you lose a few to a hyena or lion, that is bad luck.”

## Discussion

Ensuring the sustainability of protected areas for biodiversity conservation is a global issue that requires urgent attention for better conservation outcomes [1, 51]. To our knowledge, this study is the first to report on the proximate causes of illicit livestock grazing in SNP and the perception of the risks that such grazing entails. Our findings indicate that the respondents perceived that there was insufficient pasturage and water for cattle throughout the year in their areas of stay, and the number of cattle owned significantly influenced the results. Therefore, insufficient pasturage and water for cattle, and the large number of cattle owned, are perceived by respondents as the proximate causes for livestock transgression into SNP for supplementary forage. As a result, most Large Herders and Very Large Herders admitted to taking cattle into the park to supplement forage and water. In fact, most Large Herders and Very Large Herders admitted that they have been arrested inside the park multiple times compared to Small Herders and Medium Herders. Furthermore, the results indicate that regardless of the number of cattle that the livestock keepers own, they all fear being arrested inside the park. However, most of Large Herders and Very Large Herders do not fear dangerous wild animals that might attack them or their cattle, as they have different mechanisms of defense.

### Inadequate pasturage and water for livestock in the western Serengeti

Most of the respondents indicated that inadequate pasturage and water for livestock keepers is the main proximate cause for grazing inside SNP, which could be explained by the following reasons. First, livestock farming practices in east Africa, including Tanzania, have an extensive free-range grazing system in communal areas that has been in existence for decades [13, 36]. Most pastoralists in Tanzania depend on natural pastures. It should be noted that livestock-keeping efficiency depends on the quantity and quality of forage [52]. Furthermore, Tanzania is impacted by climate change, especially prolonged drought, which exacerbates the decrease in the quantity and quality of pastures for livestock [53]. The pasturage is over-utilized in areas of settlement throughout the year and hence requires supplementary forage [36, 54], which might include pasturage inside the protected areas. When the amount of pasture decreases along the protected area boundary, livestock keepers expand grazing further inside [7]. Inadequate pasturage for livestock keepers can be addressed by changes in policies governing the livestock industry in Tanzania that will promote more intensive livestock keeping and pasture management that is environmentally friendly, instead of dependence on what is occurring in the environment [55].

Second, the scarcity of water for livestock has been found to be another proximate cause that influences the transgression of livestock into SNP. The Serengeti Ecosystem experiences erratic rainfall, and most of the available water sources are seasonal streams, such as the Mbalangeti, Orangi, and Grumeti; the only perennial watercourse in the entire ecosystem is the Mara River [19]. Most of these rivers are found within protected areas such as Ikorongo, Grumeti, and SNP, and some originate outside [19]. The availability of water and rainfall patterns is known to influence the movement of livestock keepers and wildlife within the Serengeti ecosystem [36]. Recently, a study by Matata, Kegamba [9] revealed how livestock keepers in Rwamchanga Village express their concern that a newly constructed electrified fence in western Serengeti between their village and Ikorongo Game Reserve will prohibit their access to the water source located inside the reserve. The Tanzania Wildlife Conservation Act No. 5 of 2009 [56] prohibits livestock and people from entering the protected areas, and this denied access makes livestock keepers in the village feel very angry that their livestock will be negatively affected [9].

Furthermore, Kideghesho and Msuya [20] suggested that the scarcity of water in the western Serengeti is in part the result of climate change. According to Estes, Kuemmerle [25], shortages of land and water influence farmers and livestock keepers to extend grazing to protected areas. Similarly, Waweru and Oleleboo [57], in a study conducted in Tsavo West National Park in the Coast province of Kenya, noted that livestock are kept by communities close to protected areas in villages with small land and water supply. During our interviews, we found that in the dry season, livestock keepers, especially those with large herds of cattle, face many challenges in obtaining water for cattle, including traveling a long distance to the water points. In response to this challenge, Tanzania National Parks (TANAPA) has been constructing water dams in some of the villages located near the parks, but due to limited budget, only a few villages have received funding [58, 59]. We recommend that SNP management should focus on increasing the budget to support livestock keepers to build water dams for livestock on village land.

Third, the results indicate that the number of cattle owned by a respondent is another proximate cause that influences the respondents toward grazing inside protected areas. This is because most Large Herders and Very Large Herders admitted that they have been taking cattle inside the park to supplement their forage, that they have been arrested with livestock inside the park, and have been arrested multiple times compared to Small Herders and Medium Herders. The free-range grazing system forces livestock keepers to compete on common pastures, and the herders with a large number of cattle are always victorious [60]. In fact, due to the large number of cattle on common grazing land in the village and district, the amount and quality of pasturage continues to diminish due to intense competition [61]. The number of livestock appears to exceed the carrying capacity of the land, resulting in increased grazing pressure and insufficient pasture on limited land [60]. This is influenced by the land tenure system in Tanzania through the Village Land Act of 1999 (URT, 2001), which gives communities the right to use unoccupied village land as communal property [62]. Consequently, the system offers little or no protection to communal pasturage, which becomes accessible by all members of the village [63]. Therefore, livestock keepers in western Serengeti suffer Hardin’s (1968) “tragedy of the common” [64], and one of their few remaining options is supplementary forage from protected areas [8-10]. The results also indicate that for the majority of respondents in the Tarime and Serengeti districts of the Mara Region, the pasturage is even less adequate compared to that of other districts bordering SNP. This is supported by the 2019/2020 National Sample Census of Agriculture which indicates that the Mara Region is among the five leading regions in Tanzania with the highest livestock population [28].

Finally, in addition to the proximate causes identified in this study, the drivers of the root causes, specifically the lack of land use plans at the village level, can be viewed not only as the fundamental causes of livestock transgression into SNP, but also as the main source of human-wildlife conflict in the area [65]. It has been suggested that the challenge can be resolved by implementing proper land use planning for livestock on village lands adjacent to protected areas [66]. Local-level land use planning has been instrumental in solving environmental management challenges in Africa, including many land disputes since the 1980s [67, 68]. However, interviews with respondents in the surveyed villages revealed that there is no land set aside for livestock grazing, and therefore, cultivation, settlements, and livestock grazing occur within the same areas. Field observation in Kunzugu, Nyamatoke, and Machochwe villages revealed that some of the livestock keepers have already established permanent settlements and cultivation in areas that were initially set aside for livestock grazing. Most of the villages surveyed did not have land use plans; most respondents in those few villages that did, indicated that the land use plans are not respected or not implemented. Furthermore, the lack of land use plans coupled with the growing population of humans and their accompanying livestock in the villages bordering the western SNP was found to be the leading factor in encroachment onto the park boundary [25].

### Perception of the risks of grazing livestock inside the park

The results indicate that regardless of the number of cattle owned, most livestock keepers fear being arrested within the park. Their perception might be associated with historic traumatic top-down antipoaching law enforcement strategies that aim to maintain wildlife in protected areas through fines and prison sentences given to offenders [35]. Under the Wildlife Conservation Act [56] and the National Park legislation [69], entry into a protected area without a permit is deemed illegal, and offenders are liable to fines and/or imprisonment. In addition to this, some protected areas in Tanzania have reported that arrested offenders have been harassed, beaten, or had their property (cattle) confiscated by the court [70]. Therefore, to avoid being detected by wildlife rangers, most of the respondents, especially those who grazed their cattle deeper inside the park, explained that they do so at night when the movements of rangers are infrequent. Those who attempt to graze livestock inside the park during the day do so along the protected area border line, where they can easily run out to the village with their cattle when they see rangers approaching. But along the park boundary, there is not enough pasture because it is overgrazed. Our findings align with those of [35], who reported that most of the offenders in the western Serengeti conduct their grazing at night because of fear of being arrested during the day. Additionally, a recent study by Musika, Wakibara [7] indicated that the hotspots for livestock grazing in the Moyowosi Kigosi Game Reserve in Tanzania’s Shinyanga and Tabora regions are located along the boundary line.

Apart from fear of being arrested, most livestock keepers also fear dangerous animals that can harm them or their livestock while inside the park. Respondents explained that the risks of dangerous animals increase as they move deeper into the park. Dangerous animals mentioned by respondents as attacking humans and livestock include lion (*Panthera leo*), hyena (*<u>Crocuta crocuta</u>*), leopard (*<u>Panthera pardus</u>*), elephant (*Loxodonta africana*), Nile crocodile (*Crocodylus niloticus*), and African buffalo (*Syncerus caffer*). Some respondents mentioned that they tend to run away when threatened by dangerous animals, especially lions, and therefore face a great risk of losing cattle to the predators. The same risk was found in livestock keepers around Ruaha National Park in central Tanzania [6] and Moyowosi Kigosi Game Reserve in north-west Tanzania [7]. Our results are also similar to those of Butt [13], who found that the risks of livestock keepers encountering dangerous wild animals increase as they move deeper into a protected area. While hyenas, lions, and leopards lead in livestock predation in western Serengeti, locals, especially women, who collect non-timber forest products—such as building materials, fodder, and firewood—are also at great risk of encountering dangerous animals, as is shown by the human-elephant conflict in Rajiji National Park in India as humans move deeper in unfamiliar areas of the park [71].

However, most Large Herders and Very Large Herders do not fear dangerous wild animals as much, because they have access to different defense mechanisms. Respondents explained that they use different defense strategies, including powerful light torches to illuminate the area and scare carnivores, and the use of domestic dogs that help detect the presence of dangerous animals nearby. In addition, livestock keepers use traditional techniques and weapons such as a machete (*panga*), bow and arrow, and spears to defend their cattle against attacks by dangerous animals. In the Serengeti Ecosystem retaliatory killing by livestock keepers has been implicated in the extirpation of African wild dogs (*Lycaon pictus*) [72, 73].

## Conclusions

The Serengeti-Mara Ecosystem (Serengeti Ecosystem) is a major tourist destination in East Africa and offers diverse and spectacular wildlife protection. This study showed that in the face of a shortage of pasturage and water for their livestock, the majority of livestock keepers living around the western part of Serengeti National Park (SNP), perceive that there are benefits that outweigh the risk of grazing in the park. The availability of pasturage and water in the livestock keeper’s area of stay and the number of cattle owned were perceived as the main proximate causes for their grazing of livestock inside SNP. Additionally, ownership of a large number of cattle was a predominant factor for grazing of livestock in the park. Most livestock keepers who own a substantial number of cattle (Large Herders and Very Large Herders) admitted to grazing their cattle inside the SNP and have been arrested more often and more frequently compared to those with a small number of cattle (Small Herders and Medium Herders). In addition to this, inadequate pasturage and water for livestock in western Serengeti could be influenced by the extensive free-range grazing system that forces competition for livestock on a small amount of communal pasture. Although this research did not focus on the carrying capacity of the villages’ land to support livestock, the prevailing situation in the area indicates that the livestock has exceeded the carrying capacity, which has resulted in overgrazing on the community land. Moreover, in villages close to SNP, land use plans are lacking or not enforced. We recommend that the responsible authorities consider reviewing the existing policy and promote more intensive livestock husbandry, which encourages pasture management and is environmentally friendly. We also recommend that land use planning be emphasized at the village level and that strategies be adopted that can ensure its sustainable implementation. Furthermore, global climate change could be one of the factors behind the scarcity of water for livestock keepers in the western Serengeti. We also recommend that Serengeti National Park (SNP) continues to build farm dams and ponds on village land to increase retention by catchment and retention of rainy season water for livestock keepers.

Furthermore, we found that livestock farmers in the western Serengeti take significant risks in grazing livestock in the park. Common risks the respondents mentioned include being harassed by wildlife rangers after being arrested, beatings, fines, imprisonment, confiscation of cattle by the court, loss of cattle in the wilderness, and the threat of attack from dangerous wild animals. However, regardless of those risks and the number of cattle owned, the majority of respondents fear being arrested in the park. Furthermore, the majority of respondents fear dangerous animals that may attack them or their cattle in the park, although most of the respondents with a significant number of cattle have less fear of dangerous animals because they have access to more defense mechanisms compared to those with a smaller number of cattle.

## 1. Acknowledgements

The authors thank the District Executive Directors of Serengeti, Serengeti, Bariadi, Simiyu, Tarime and Bunda districts for granting permits to collect data and their cooperation. We are grateful to the community respondents, the community leaders, and the residents of the surveyed villages who provided unequivocal support during the surveys. The authors thank the College of African Wildlife Management, Mweka, for the field financial support that made this work achievable. We also thank the authorities of Serengeti National Park specifically, Mr. Stephano Kimera Msumi (Head Law Enforcement Department, during this survey), for his cooperation. Finally, we thank Mr. Leo Atwood for technical English editing.

## Ethics approval and consent

The College of African Wildlife Management, Mweka through the Research Department provided a research clearance including ethical issues for consideration during the entire research. The office of District Executive Director was officially informed about the research before data collection. Oral consents were also obtained from each respondent before interview.

## S1. Appendix: Questionnaire for household survey

Dear participant,

My name is Maria Lameck E. Matungwa / Juma J. Kegamba, from the College of African Wildlife Management Mweka. I am doing a research on **assessment of livestock grazing pressure in the western part of Serengeti National Park**. I am asking for your attention and cooperation in answering the following questions and the information obtained will be used for study purpose and not otherwise.

After reading this information/been informed, are you ready to grant your permission for the use of your statement/words for this research?

## A. Respondent’s demographic information

1. Village Name………………………………….. District…………………….
  a. Gender
  b. Male ()
  c. Female ()
2. Age
  a. 18 – 35 years ()
  b. 36 – 55 years ()
  c. Above 55 years ()
3. How many cattle do you have
  a. 1-20
  b. 21-50 ()
  c. 51-100 ()
  d. >100 ()
4. Residence status
  a. Born in the village ()
  b. Internal migrant ()

## B. The knowledge of the respondent about the availability of pastures and water for livestock in their area of stay (district)

1. Do you have specific area in the district set aside for livestock grazing? Yes () No ()
2. If “Yes” in 6 above do the grazing areas you have in your area enough for grazing livestock throughout the year? Yes () No ()
3. Are the water for livestock in your areas enough throughout the year? Yes () No ()
4. If “No” in 8 above where do you normally take your livestock to complement the scarcity of grazing pastures and water? …………………………………………………………

## C. The experience of the respondents about grazing cattle inside SNP

1. Have you ever taken cattle to graze in the park? Yes () No ()
2. Have you ever been arrested inside the park for grazing cattle? Yes () No ()
3. If “Yes” in 11 above, how many times you have been arrested None (), Once () and multiple times ()
4. Explain what happened after you were arrested……………………………………… …………………………………………………………………………………………….

## D. Respondents fear of being arrested by rangers or fear of problematic wild animals in SNP

1. Do you fear of being arrested inside the park? Yes () No ()
2. If Yes explain why do you fear…………………………………………………………
3. Do you fear the risks of dangerous wild animals that may attack you or your cattle inside the park? Yes () No ()
4. If Yes explain why …………………………………………………………
5. If No explain why …………………………………………………………
6. If “No” in 20 above do you face any problem when taking your livestock for grazing in SNP? Yes () No ()
7. What do you think should be done to resolve pasture and water challenge for livestock keeping in your areas?……………………………………………………………… ………………………………………………………………………………………………

